# RNA self-association limits the removal of double-stranded RNA by affinity chromatography

**DOI:** 10.64898/2026.09.21.749193

**Authors:** Nathaniel E Clark, Heather Mallory, Christian J Ndong, Matthew R Schraut, Roger A Winters, Jennifer Freise, Brandon L Kier, Matthew J Ranaghan, Lisa Deprez, Senne Dillen, Kelley Kearns, Jing Zhu, Thomas C Scanlon

## Abstract

Double-stranded RNAs are inflammatory byproducts of *in vitro* transcription of single-stranded RNA. Here we investigate removal of dsRNA byproducts using dsRNA affinity chromatography (dsRNA-AC) and two common mRNA molecules: 1) GFP, and 2) high dsRNA wild-type firefly luciferase (fLuc_WT_). For GFP mRNA, dsRNA levels decreased by 2,000-fold, and mRNA recovery was high. In contrast, dsRNA levels decreased only two-fold for fLuc_WT_, and mRNA yields were lower. Biophysical characterization revealed that fLuc_WT_ contains polydisperse higher-order structures that interfered with dsRNA-AC. Sedimentation-velocity analytical ultracentrifugation experiments demonstrated the higher-order structures were driven by mRNA concentration. Decreasing fLuc_WT_ concentrations increased the effectiveness of dsRNA-AC by reducing polydispersity. dsRNA levels were measured with ELISA, dot-blots, immuno-northern blots, and a cell-based interferon release assay, enabling comparison and cross-validation of these analytical methods. dsRNA-AC is a new chromatography modality, and this study identifies sample polydispersity as a key determinant of efficient dsRNA removal.

## Introduction

Single-stranded RNA therapeutics such as messenger RNA (mRNA), self-amplifying RNA (saRNA), and circular (circRNA) represent a new platform technology for vaccines and gene therapies (1,13,15). Despite rapid clinical success in response to the COVD-19 pandemic, significant manufacturing challenges remain, such as the production of double-stranded RNA (dsRNA) impurities during in-vitro transcription (2,5,9,17). dsRNA activates innate immune pathways, including RIG-I and Toll-like receptors, leading to reduced translation efficiency and increased inflammatory responses, which can manifest as flu-like side effects.

Traditional purification approaches such as poly deoxythymidine (dT) chromatography and tangential flow filtration do not effectively remove dsRNA due to similar physicochemical properties of the dsRNAs to the target molecules (10). Alternative methods such as reversed-phase chromatography and cellulose purification reduce dsRNA but face limitations in scalability and process compatibility due to organic solvent requirements (6,16,18). Affinity chromatography offers a targeted approach based on dsRNA-specific recognition. Because dsRNA-AC is an aqueous operation, it resembles traditional, scalable bioprocess operations such as those used for monoclonal antibodies and protein therapeutics.

Previously, we examined the ability of dsRNA-AC to reduce the immunogenicity of three different preparations of GFP mRNA(7). Here, we test dsRNA-AC using an additional mRNA: wild-type firefly luciferase (1.8 kb, fLuc_WT_). We evaluated the effect of the following factors on dsRNA removal efficiency: mRNA concentration, salt type, and salt concentration. dsRNA levels were analyzed with immuno-analytical methods such as ELISA, dot-blots, and immuno-northern blots (INB), and a cell-based interferon release assay. Structural differences between the mRNA samples were investigated with circular dichroism, size-exclusion chromatography, mass-photometry, dynamic light scattering, and sedimentation velocity analytical ultracentrifugation.

## Results

### Batch-binding experiments

Initially we tested dsRNA-AC in small-scale batch binding experiments (Figure 1A). A total of 2 samples were tested; a lithium chloride precipitate of fLuc_WT_, and a commercial GFP preparation. Samples were constituted in 20 mM Tris pH 7.5, 1 mM EDTA, and 0.63 M NaCl at mRNA concentrations of 0.46 (GFP) and 0.64 (fLuc_WT_) mg/mL. The experimental parameters (feed titer and resin challenge) and results (dsRNA levels, mRNA yield, and mass balance) are summarized in Figure 1B.

**Figure 1.**
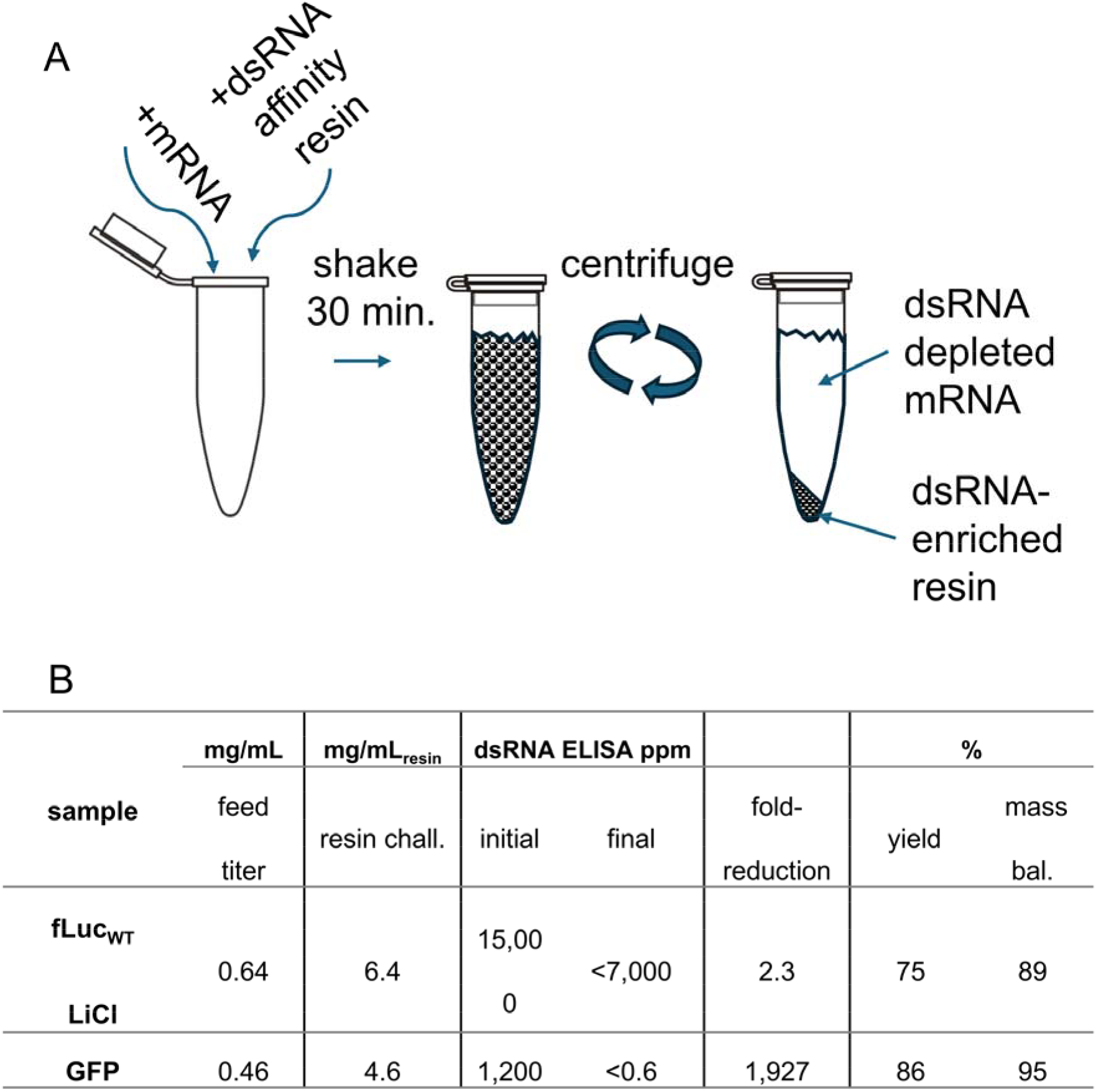
Overview of batch binding experiment. A)mRNA samples are incubated with a dsRNA-selective affinity resin. After a 30-minute incubation, the resin is separated by centrifugation from the dsRNA-depleted mRNA in the supernatant. The dsRNA-enriched material is bound to the resin. B) Table of experimental parameters and results

For GFP, dsRNA levels decreased from 1,200 to <0.6 ppm, a 1,927-fold reduction. For fLuc_WT_, dsRNA decreased from 15,000 to <7,000 ppm, a modest 2.3-fold reduction. The mRNA yields were 86% for GFP, and 75% for fLuc_WT_ (Figure 1B)

### RNA solubility time course assays

To test if the solubility of these molecules differed in the high-salt purification buffer, we prepared samples in water, 0.63 M NaCl, or 0.63 M KCl (each with 20 mM Tris pH 7.5 and 1 mM EDTA). The samples were incubated at 20° C on an orbital mixer, and at the indicated timepoints the samples were centrifuged and soluble RNA titers measured (Figure 2).

**Figure 2.**
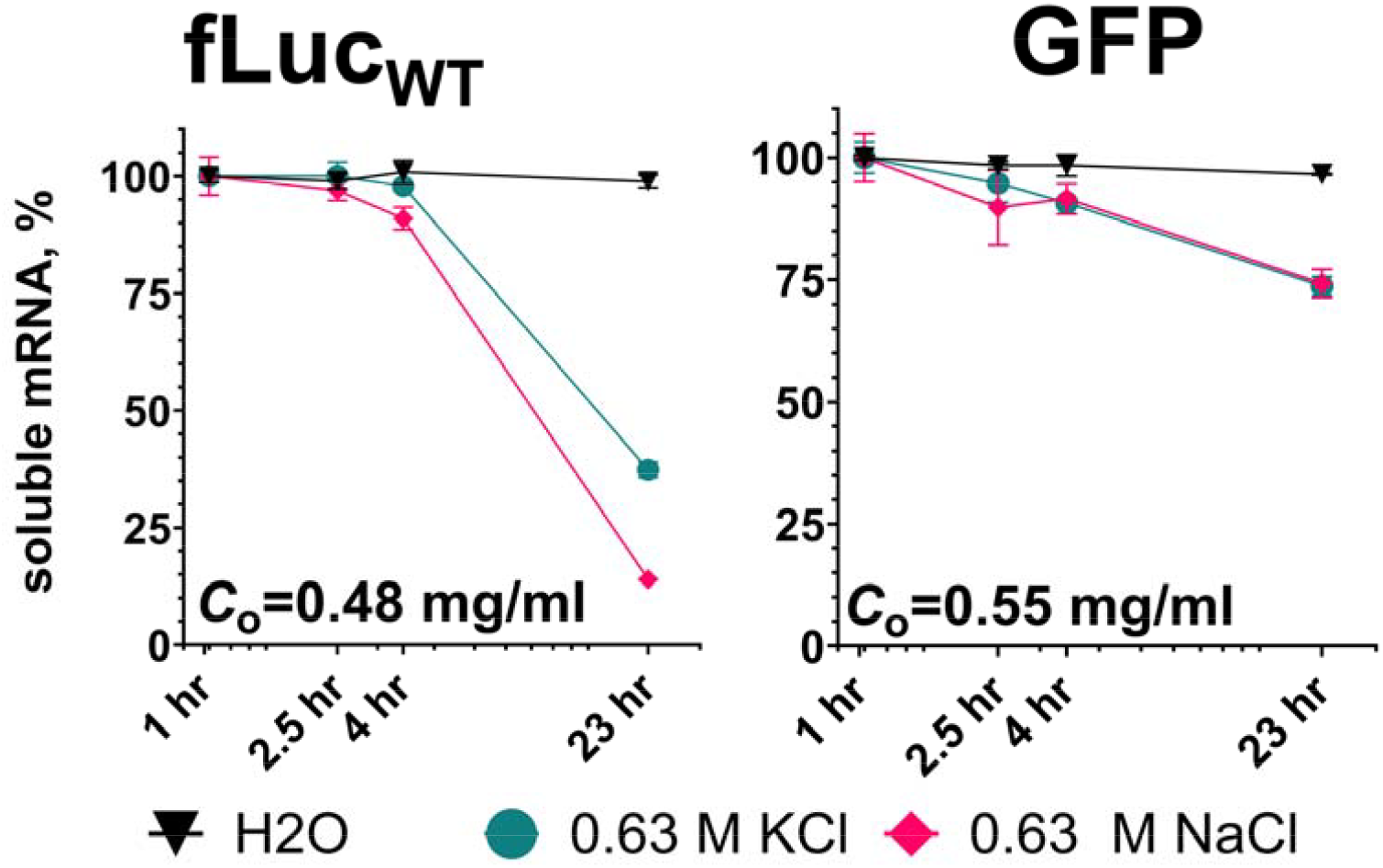
Time course of mRNA solubility in NaCl and KCl. Each mRNA is constituted in water, 0.63 M NaCl, or 0.63 M KCl and incubated at room temperature with shaking. At the indicated times, samples were centrifuged to separate precipitated and soluble mRNA. Soluble mRNA concentrations were measured with spectrophotometry and plotted as % of initial mRNA concentration. The initial mRNA concentrations (*C*_o_) are indicated above the x-axes.

In water, both RNAs remained soluble for the entire 23-hour experiment. In NaCl, fLuc_WT_ was relatively soluble between 2.5 and 4 hours, but >70% precipitated at 23 hours. GFP was highly soluble in NaCl and KCl for 23 hours. fLuc_WT_ solubility was better in KCl, with slower time-dependent precipitation for fLuc_WT_.

### Biophysical characterization of the mRNA feeds

We hypothesized that the divergent results in the solubility and batch-purification assays were due to structural and biophysical differences intrinsic to two distinct primary sequences. We employed an suite of biophysical techniques to characterize the secondary structure, size distribution, and aggregation state of each mRNA construct in solution.

We collected circular-dichroism (CD) spectroscopy to quantify differences in secondary structure of each mRNA. Spectra collected from 10 to 95 °C were broadly similar between the two mRNAs (Figure S2A). Analysis of the 263 nm peak of normalized spectra revealed that fLuc_WT_ melted at 45 °C, while GFP melted around 60 °C (Figure S2B). Normalized spectra and other CD metrics showed no major differences between samples (Figure S2B)

The peak width of the mRNAs in NaCl tracked with the results from the batch binding (Figure 1B) and solubility time course (Figure 2) assays. In the sample with a narrower SEC peak width (GFP, 0.5 mL), dsRNA reduction (Figure 1B) and NaCl solubility (Figure 2) were higher. In the sample with a wider SEC peak (fLuc_WT_, 2.4 mL), dsRNA reduction (Figure 1B) and NaCl solubility (Figure 2) were lower.

Mass photometry (MP) was used to quantify the sizes of the various mRNA species in solution (3,11). Samples were prepared in PBS for optimal surface adsorption (4). Both RNAs presented mass histograms with a dominant species that measured within approximately 2% of their calculated sizes (Figure 3B, Table 1). A small dimer peak was observed for GFP. fLuc_WT_ showed large peaks for dimer and higher-order structures, consistent with corresponding SEC profile in water (Figure 3A).

**Table 1.**
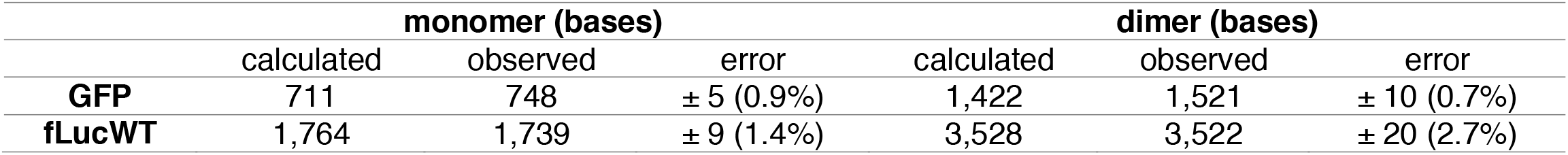
mRNA sizes by mass photometry.

|  | monomer (bases) |  |  | dimer (bases) |  |  |
| --- | --- | --- | --- | --- | --- | --- |
|  | calculated | observed | error | calculated | observed | error |
| **GFP** | 711 | 748 | ± 5 (0.9%) | 1,422 | 1,521 | ± 10 (0.7%) |
| **fLucWT** | 1,764 | 1,739 | ± 9 (1.4%) | 3,528 | 3,522 | ± 20 (2.7%) |

**Figure 3.**
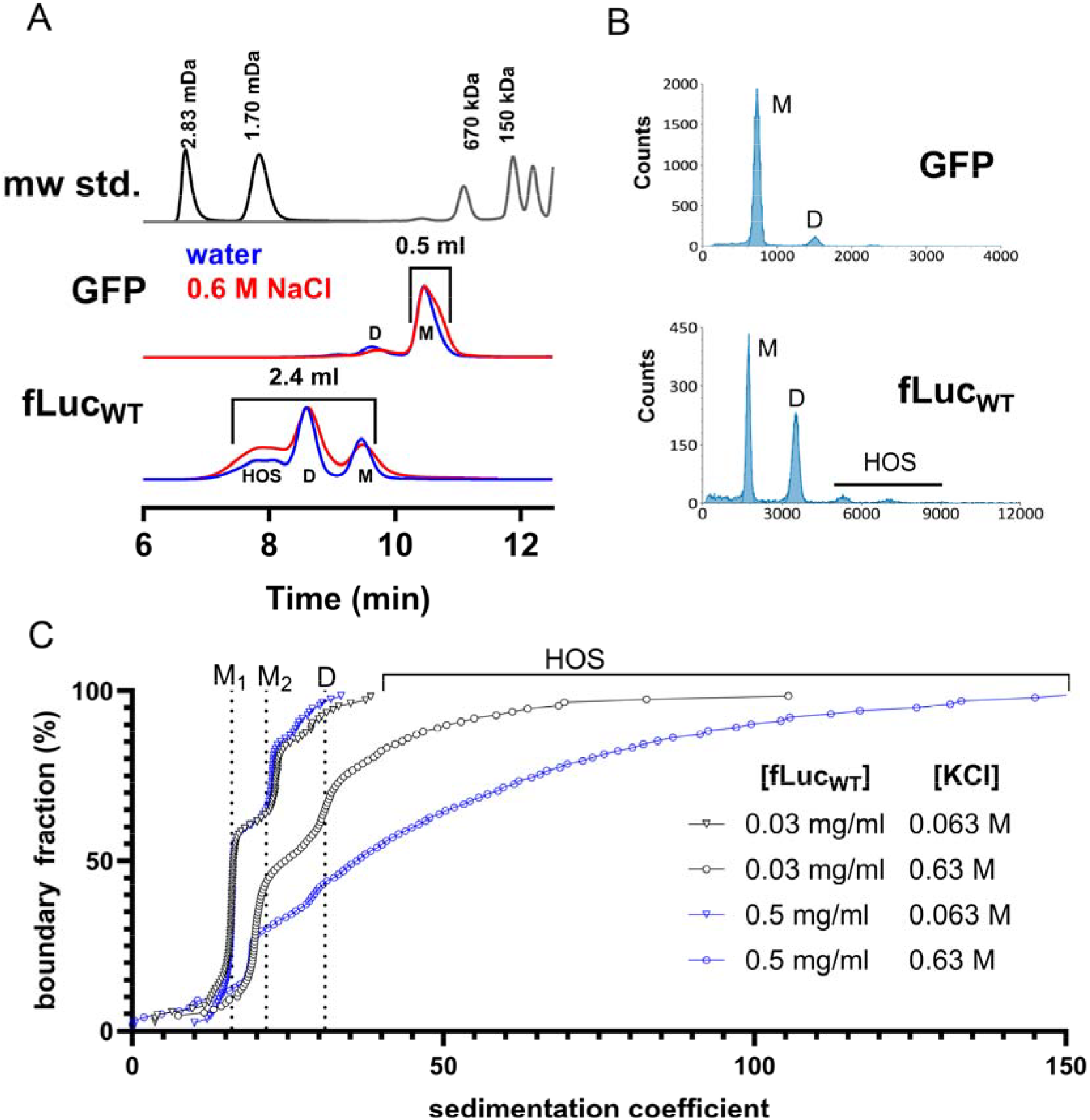
Biophysical characterization of the mRNAs. (A) Normalized chromatograms (wide-pore 1,000 Å SEC column) of mRNAs incubated in water, or 0.63 M NaCl, 20 mM Tris pH 7.5, 1 mM EDTA. (B) MP histograms of mRNAs (5 nM) in PBS. (A-B) M=monomer, D=dimer, HOS=higher-order structures. (C) Distribution of sedimentation coefficients from SV-AUC experiment. The mRNA and KCl concentrations are inset. M_1_=extended monomer, M_2_=compact monomer, D=dimer, HOS=higher-order structures

To measure the polydispersity of fLuc_WT_ at the salt and mRNA concentration used for dsRNA-AC processing, we performed sedimentation-velocity analytical ultracentrifugation (SV-AUC) experiments. To determine if formation of dimers and higher-order structures was driven by the mRNA concentration, we prepared samples at 0.5 or 0.03 mg/mL RNA in either 0.63 KCl or 0.063 M KCl. In low salt, the distribution of sedimentation coefficients was similar at both RNA concentrations (Figure 3C). Two species were observed, an extended monomer at 16 s (M_1_), and a compact monomer at 21 s (M_2_). These assignments were based on expected values for similarly sized RNAs (20). In high salt, the extended monomer was not observed, and dimers and higher-order structures formed in an RNA concentration-dependent manner. At 0.03 mg/mL fLuc_WT_, the sample contained approximately 50% compact monomer, 25% dimer, and 25% higher-order species. At 0.5 mg/mL, the distribution shifted to approximately 30% compact monomer, 10% dimer, and 60% higher-order species (Figure 3C).

As a result of these experiments, we concluded that fLuc_WT_ formed concentration-dependent higher-order structures in the presence of 0.63 M salt. With this understanding, we proceeded to purify this RNA with dsRNA-AC column runs.

### Column runs with fLuc_WT_

At feed titers of ∼0.5 mg/mL, fLuc_WT_ column runs produced typical flow-through and elution peaks (Figure 4; Table 2). In NaCl, dsRNA decreased from 1.5% to 0.6%, with 78% yield and 94% mass balance. In KCl, dsRNA decreased from 1.9% to 1.2%, with 92% yield and 96% mass balance.

**Table 2.**
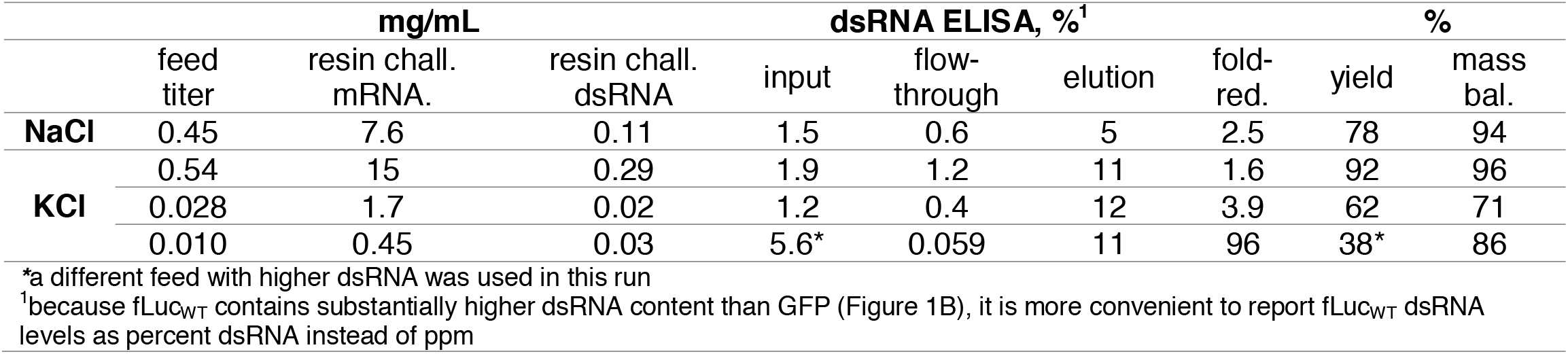
Column runs with fLuc_WT_ mRNA.

**Figure 4.**
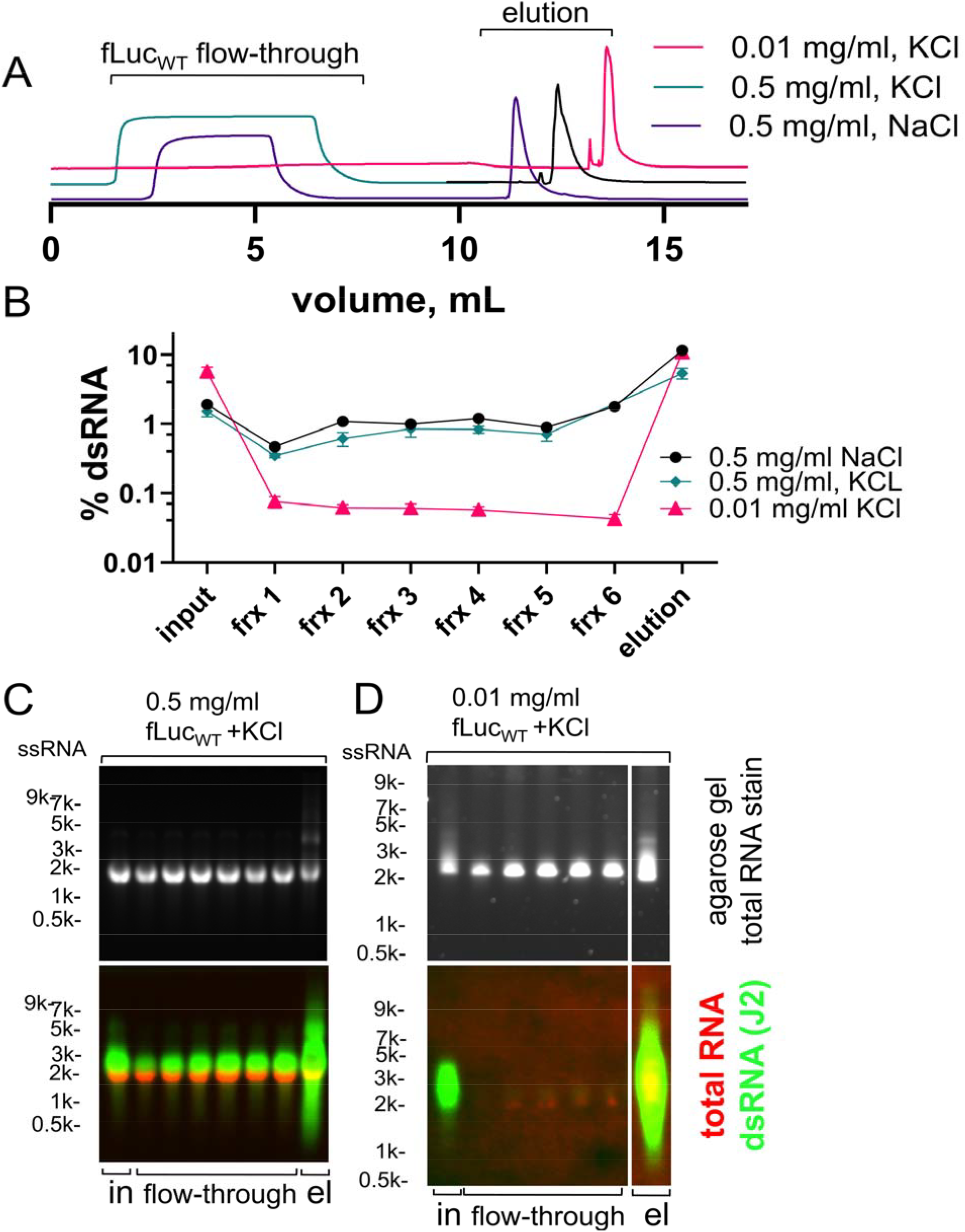
Purification of fLuc_WT_ with dsRNA affinity chromatography. (A) Offset dsRNA-AC traces for the 4 purification runs. (B) Quantification of the dsRNA levels in the process samples with ELISA. (C) Agarose gel of the 0.5 mg/mL KCl column run (top), and J2 immuno-northern blot (bottom). (D) Agarose gel of the 0.01 mg/mL KCl column run (top), and J2 immuno-northern blot (bottom).

fLuc_WT_ displayed the expected input/flow-through/elution partitioning of dsRNA by both ELISA and immuno-northern blot (Figure 4B, C, Table 2), where the input has more dsRNA than the flow-through, and the elution is highly enriched in dsRNA. However, dsRNA levels decreased by only ∼50%, and the dsRNA content of the flow-through remained very high, with results from both ELISA (Figure 4B) and immuno-northern blots (Figure 4C) in agreement on this point. The total challenge of dsRNA (0.1-0.3 mg dsRNA/mL_resin_, Table 2) was not in excess of the stated capacity of the affinity resin (0.4 mg dsRNA/mL_resin_). Therefore, we reasoned that either the effective capacity was lower for the specific fLuc_WT_ dsRNA molecules, or the self-associative behavior of fLuc_WT_ at 0.5 mg/mL interfered with dsRNA capture.

To determine if fLuc_WT_ self-association interfered with dsRNA capture, we performed additional column runs with lower feed titers. At 0.028 mg/mL, the dsRNA content of the flow-through was 0.4%. Further decreasing the feed titer to 0.010 mg/mL resulted in a flow-through dsRNA content of 0.06% (Table 2, Figure 4B), representing a 2-log reduction of dsRNA content. The immuno-northern blot detected no dsRNA in the flow through fractions of the 0.010 mg/mL run (Figure 4D). These observations suggested that self-association was the cause of low dsRNA removal, but due to the dilute feed, we were unable to load the column to the point of dsRNA breakthrough. Therefore, we could not rule the possibility of very low capacity for the fLuc_WT_ specific dsRNA molecules.

To determine if the effective capacity for fLuc_WT_ dsRNA was lower than the specified capacity, we performed column runs with two different dsRNA molecules, a purified 500 base-pair dsRNA, and an 1,800 base-pair duplex of fLuc_WT_ (Figure 5A). The dynamic binding capacity at 10% breakthrough (DBC_10_) for the 500 bp dsRNA was 0.4 g/L, consistent with the product specifications. The DBC_10_ for the 1,800 bp fLuc_WT_ duplex was slightly lower at 0.31 g/L. This decrease was insufficient to explain the high dsRNA breakthrough in column runs with 0.11 and 0.29 g/L_resin_ dsRNA challenges (Figure 4B,C, Table 2). This finding suggests the lack of dsRNA removal is due to fLuc_WT_ self-association, and not column capacity.

**Figure 5.**
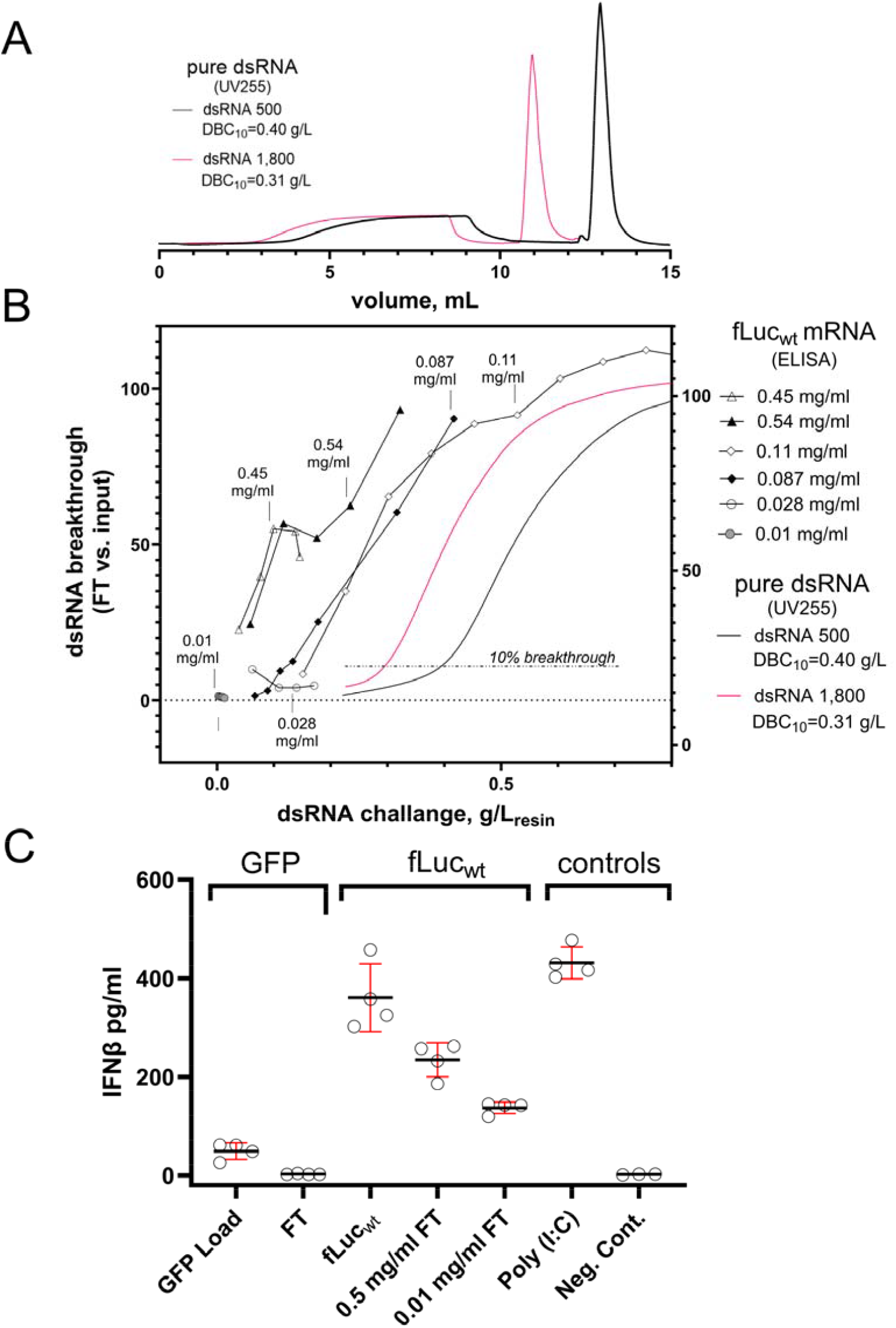
dsRNA breakthrough curves and cell-based assays. (A) Chromatograms of a 0.2 mL dsRNA affinity column loaded to breakthrough with 500 or 1,800 bp dsRNAs. (B) Plot of dsRNA breakthrough versus dsRNA challenge for 6 fLuc_WT_ runs, and the 500 and 1,800 bp dsRNA runs. dsRNA levels were quantified with ELISA. Feed titers are listed for each fLuc_WT_ run, and the 10% breakthrough values are listed for the 500 and 1,800 bp dsRNAs. (C) Cell-based interferon release assay. A549 cells were transfected with the indicated mRNA or control treatment, and the IFNβ levels in the culture media were quantified with an ELISA. Each condition was tested in quadruplicate, and the error bars represent 1 standard deviation.

Plotting the ELISA data from 6 different fLuc_WT_ column runs revealed that as fLuc_WT_ feed titer decreases from ∼0.5 to ∼0.1 mg/mL (triangles vs. squares, Figure 5B), the dsRNA breakthrough curve shifts to the right, reflecting higher effective capacity. This observation links the dsRNA clearance limitation at high fLuc_WT_ feed titers to the concentration-dependent self-association observed with SEC, mass photometry, and SV-AUC.

### Cell-based immunogenicity assay of purified RNAs

We then asked if the clearances of dsRNA observed in the various immunoassays correlated to a reduction in cell-based immunogenicity. A549 cells were transfected with mRNA pre- and post-dsRNA-AC, and interferon beta (IFNβ) response was measured with an ELISA. The positive control was poly(I:C), a toll-like receptor-3 (TLR-3) agonist, and the negative control was untreated cells (Figure 5C). These controls established the high (440 pg/mL) and low (2 pg/mL) limits of the assay. GFP transfection generated 50 pg/mL IFNβ pre-dsRNA-AC, and baseline levels of ∼2 pg/mL post dsRNA-AC. fLuc_WT_ transfection induced 360 pg/mL. dsRNA-AC purification reduced the IFNβ to 260 pg/mL (0.5 mg/mL column run), and 140 pg/mL (0.01 mg/mL column run). This was consistent with the trend of higher dsRNA removal with lower feed titer for fLuc_WT_. The relatively high residual immunogenicity may be due to an inefficient post-transcriptional capping reaction.

## Discussion

This study shows that dsRNA affinity chromatography can selectively remove immunostimulatory dsRNA from mRNA preparations. The two mRNAs tested differ in sequence, length, and dsRNA content. We summarized the challenges and solutions for each mRNA (Table 3). GFP was a favorable substrate, with strong dsRNA depletion, high mRNA recovery.

**Table 3.** Summary of experimental results.

| mRNA | analytical issues? | self-association? | dsRNA reduction?<br>(by assay) |  |  | solution |
| --- | --- | --- | --- | --- | --- | --- |
| | | | immuno-northern blot | ELISA | IFNβ assay | |
| fLuc<sub>WT</sub> | no | significant | yes | yes | yes | decrease feed titer to decrease self-association |
| GFP | no | no | yes | yes | yes | not needed |

In contrast, fLuc_WT_ presented challenges during the dsRNA-AC process. SV-AUC revealed that fLuc_WT_ formed higher-order structures in high salt. Formation of these large species was RNA concentration-dependent. These higher-order structures reduced the effective capacity of the column for dsRNA. This cause was somewhat difficult to pinpoint, because the higher-order species remained soluble through dsRNA-AC operations. Despite the ∼50% of the fLuc_WT_ sedimenting at s-values of 30-150 in high-salt, these self-associated fLuc_WT_ molecules remained in solution, and mRNA yields were relatively high in the batch and column purifications (Figure 1B, Table 2). The salt solubility time course revealed a relatively slow rate of precipitation during the first 4 hours (Figure 2). The slow rate of salt-induced precipitation appears to be a unique property of RNA solutions. In contrast, protein solutions precipitate rapidly and immediately in salting-out experiments.

The series of biophysical experiments highlighted the fLuc_WT_ higher order structures (HOS) as the most significant difference between the mRNAs. Only fLuc_WT_ had significant HOS in SEC runs without salt (Figure 3A), and was the only sample where significant dimers and HOS were detected with mass photometry (Figure 3B). SV-AUC confirmed that HOS were driven by self-association in high salt. As to precisely how the presence of HOS interferes with dsRNA-AC, we propose two possible mechanisms: i) sequestration of the dsRNAs in higher-order structures prevents capture by the affinity ligand, ii) pseudo-dsRNA epitopes present in higher order structures compete with true dsRNA for binding sites on the affinity ligand. Both mechanisms appear consistent with our experimental results.

We observe that the elution fraction for fLuc_WT_ contains a portion of mRNA that is trapped in HOS which can be removed by heat treatment (Figure S5). These species could contain pseudo-dsRNA epitopes which may interact with the ligand, or simply be precipitates that are trapped by the column frit and resolubilized (but not completely denatured) in elution buffer. In either case, it appears that the column is removing these HOS from the flow-through fractions. This suggests that the yield of RNA in the flow-through could increase if these HOS are controlled or reduced before the dsRNA-AC operation.

We find that low polydispersity is correlated with the efficient dsRNA-AC. However polydispersity is not typically measured and controlled during RNA purification development. As RNA therapies become more common and manufacturing processes more refined, we believe that tools for measuring RNA polydispersity will be valuable. SEC provided some valuable information, but the method is non-equilibrium. Sample polydispersity can change during the run due to differences between the sample buffer and mobile phase, and dissociation driven by shifting mRNA concentrations. MP provided a useful indicator of problematic self-association, as only fLuc_WT_ had significant dimer and HOS in the low salt analysis buffer (Figure 3B). We were unable to measure at high salt due to poor adsorption to the glass slide, but this issue can in principle be overcome with advances in the surface capture chemistry. SV-AUC was very well suited to measuring the structural complexity of fLuc_WT_ in the exact salt and mRNA concentrations of a dsRNA-AC process. The UV optics enabled high-sensitivity detection of samples spanning a 17-fold range of concentrations, and the experiment was compatible with the 0.63 M KCl condition. Thus, SV-AUC is a powerful method to analyze RNA dispersity in bioprocessing-relevant conditions.

This study demonstrates that dsRNA affinity chromatography selectively removes immunostimulatory dsRNA from RNA preparations. Purification success is primarily determined by the mono/polydispersity of the sample in the processing condition, a sequence-intrinsic property. Identifying appropriate analytical methods for RNA polydispersity enabled optimization of the dsRNA-AC process through control of higher order structures. fLuc_WT_ serves as a useful extreme example of a sequence with high propensity to self-associate. Effective dsRNA removal required a very low mRNA titer, which would result in excessive buffer usage and sample volumes at manufacturing scale. A better solution would be sequence optimization with the goal of reducing self-association. Sequence optimizations typically aim to increase translation, lower/control the guanine and cytosine (GC) content, and decrease uridine (U) content (8,14). In addition to the many constraints on RNA sequence design, sequences with low polydispersity (e.g., GFP) perform better in downstream processing. Therefore, sequence design presents an opportunity to improve manufacturing efficiency and economics by controlling polydispersity of the target RNA.

RNA concentration emerged as a critical process parameter. For many RNAs, dsRNA-AC is a simple high-yield operation. We have encountered a subset of RNAs for which dsRNA-AC presents various challenges, and these are the RNAs which self-associate at high concentrations, leading to analytical artifacts, and/or decreased effective resin capacity. fLuc_WT_ represents the most extreme example of a self-associating RNA that we have encountered. These findings indicate that process optimization must consider the unique structural and biophysical properties of a specific RNA molecule to achieve consistent removal of dsRNA byproducts. Once these factors are understood for the more challenging RNAs, affinity chromatography provides a scalable and efficient strategy for improving the quality of all single-stranded RNA therapeutics, including mRNA, circRNA, and saRNA.

## Conclusion

dsRNA affinity chromatography provides a scalable strategy for removal of immunostimulatory dsRNA impurities. Performance depends on maintaining relatively monodisperse RNA samples, with minimal quantities of higher-order structures. Dispersity is primarily determined by RNA sequence, RNA concentration, and buffer ionic strength. Not all RNAs have the same structural and biophysical properties, and processing conditions (salt and RNA concentration in particular) must be tailored for the individual RNA, based on the results of hydrodynamic analyses such as SV-AUC. Orthogonal dsRNA analytical approaches enable confident assessment of dsRNA levels and clearance of immunogens during dsRNA-AC process development.

## Materials and Methods

### mRNA sourcing and synthesis

GFP mRNA (Figure 1B, Figure 2) was purchased commercially (DasherGFP® mRNA, Aldevron 3870). For cell based assays, the optimized IVT GFP samples pre- and post-dsRNA-AC were used (7). fLuc_WT_ was prepared using a commercial IVT kit and the control template (HiScribe™ T7 High Yield RNA Synthesis Kit, New England Biolabs E2040S), capped using Vaccinia Capping Enzyme (New England Biolabs M2080) and 2’-O-methyltransferase (New England Biolabs M0366), and tailed using E. coli Poly(A) Polymerase (New England Biolabs M0276).

The 500 base-pair dsRNA was prepared using a commercial kit (MEGAScript RNAi kit, Thermo Fisher Scientific AM1626), then purified by dsRNA-AC to remove single-stranded RNA. Preparation of 1,800 base-pair fLuc duplex from two complementary strands was described previously (10).

### dsRNA removal by affinity chromatography

Double-stranded RNA (dsRNA) impurities were removed using a dsRNA-selective affinity resin (AVIPure® dsRNA Clear Affinity Resin, Repligen 100RNA-5). For batch binding, mRNA samples were diluted into high-salt binding buffer (20 mM Tris, 0.63 M NaCl, 1 mM EDTA, pH 7.5), and each 0.5 mL RNA sample was mixed with 50 µL of equilibrated resin, and incubated with mixing at room temperature for 30 minutes using a ThermoMixer at 700 RPMs (Eppendorf). Following incubation, resin was pelleted with a microcentrifuge, and the supernatant (FT) transferred to a new tube. Resin was washed with binding buffer, and bound material was eluted using 6 M guanidinium hydrochloride (GuHCl; Thermo 24115).

Column-based purification was performed using 0.2 mL bed-volume prepacked chromatography columns (Repligen 23052205-02) on an NGC chromatography system with multi-wavelength detector (BioRad 7880009). A buffer of 20 mM Tris, 1 mM EDTA, 0.625 M KCL or NaCl, pH 7.5, was used for sample preparation, column equilibration, and column washing, as indicated in the figures and tables. For elution, 6 M GuHCl or 0.1% SDS were used interchangeably. Both solutions reversibly unfold the affinity ligand and release bound dsRNA.

### Agarose gel electrophoresis

mRNAs were analyzed by native agarose gel electrophoresis using 1.5% agarose gels, 1x TBE running buffer, and SybrSafe stain (Thermo S33102), as previously described (7,10). ssRNA and dsRNA ladders were loaded on each gel (New England Biolabs N0362, N0363). Samples were not heated prior to electrophoresis except in Figure S5, where the samples were heated at 72□°C for 2 minutes.

### Salt-solubility assay

The indicated RNAs were prepared in either water, 20 mM Tris, 1 mM EDTA, 0.625 M KCl, pH 7.5, or 20 mM Tris, 1 mM EDTA, 0.625 M NaCl, pH 7.5. 0.5 mL samples were incubated at room temperature with mixing with a Thermomixer at 700 rpm (Eppendorf). At the indicated timepoints, the samples were centrifuged in a microcentifuge for 3 minutes at an RCF of 16,100, and the soluble RNA concentration measured with UV spectroscopy (NanoDrop Lite, Thermo ND⍰LITE⍰PR).

### dsRNA detection and quantification

dsRNA content was quantified using a sandwich enzyme-linked immunosorbent assay (EasyAna dsRNA ELISA kit, Vazyme, Cat. No. DD3509). dsRNA concentrations were calculated using a four-parameter logistic (4PL) fit. Immuno-northern blots and immuno-dot blots used HRP-conjugated J2 antibody (Novus, NBP3⍰11395H), as previously described (10).

### Cell-based interferon assay

A549 cells (ATCC, PTA-6231) were cultured in F12 medium with 10% FBS and transfected with lipid-based transfection reagent (Thermo LMRNA008) in 96-well plates. Prior to transfection, all RNAs were buffer exchanged into water using 30k-MWCO spin concentrators (Millipore UFC803024) and adjusted to 0.4 mg/mL. After 24 hours, culture media was collected, and IFNβ levels were quantified by ELISA (R&D Systems DY814-05). High-molecular weight poly(I:C) was the positive control (Invivogen tlrl-pic), and cells in media were the negative control. Each sample and control was assayed in quadruplicate. IFNβ levels were calculated using a four-parameter logistic (4PL) fit.

### Circular dichroism

The RNAs were prepared in water at 0.1 mg/mL and placed in 1 mm quartz cuvettes. Absorbance spectra were collected from 190-330 nm, with 0.1 nm resolution, 1 nm bandwidth, using a Jasco J-815 spectropolarimeter. Samples were measured at 10° C, then at 5° increments up to 95° C. Raw data were exported to Prism for plotting and analysis.

### Size Exclusion Chromatography

mRNAs were analyzed with a 1,000 Å wide-pore analytical SEC column (GTxResolve Premier SEC 1000 Å, 3 μm, 4.6 x 300 mm, Waters 186010736). The mobile phase was 0.1 M sodium phosphate, 0.2 M sodium chloride, pH 6.8. 3 µL of sample was injected, with a flow rate of 0.3 mL/min. Molecular weight standards included a DNA vector mix (2.83 mDa and 1.70 mDa peaks, Waters 186011285), and two sets of protein standards (BEH200, Waters 186006518, and AdvanceBio 130, Agilent 5190-9416). To aid in comparison, all chromatograms were normalized in figure 3A.

### Mass photometry

All MP experiments were performed using a Refeyn TwoMP instrument (Refeyn Ltd). Samples were prepared at 5 nM in 1x PBS (approximately 0.001 mg/mL) using a MassGlass NA slide (Refeyn Ltd). Movies were collected in Normal Mode with a Large Field of View for 60 seconds. RNA samples were calibrated against a single-stranded RNA ladder (New England Biolabs). Data were processed using DiscoverMP software (Refeyn Ltd). Measured RNA lengths were compared to expected transcript sizes (Supplemental File 1) to confirm sample identity (Figure 3B).

### Sedimentation-velocity analytical ultracentrifugation

fLuc_WT_ was prepared at 0.03 mg/mL and 0.5 mg/mL in 0.063M KCl, 20 mM Tris, 1 mM EDTA, pH 7.5, or 0.63M KCl, 20 mM Tris, 1 mM EDTA, pH 7.5 (4 samples total). Analysis was performed at the Northern Rockies Center for Hydrodynamics, University of Montana. The experiment was performed at 36,000 rpm, 4 ºC, for 10 hours, using an Optima AUC (Beckman). UV absorbance was measured at 299 nm for the 0.5 mg/mL samples, and 273 nm for the 0.03 mg/mL samples. Data were analyzed using two-dimensional spectrum analysis (19) and the standard analysis workflows in the UltraScan analysis software (12).

## Supporting information

Supplemental Sequences

## Data analysis and presentation

Data were plotted using GraphPad Prism, version 11 (GraphPad Software). Statistical significance was determined with an unpaired Welch’s t-test (Figure 5C). 4PL fits of ELISA data were calculated with Excel (Microsoft).

### Artificial Intelligence statement

Microsoft Copilot generated an initial draft of the manuscript. This draft was extensively rewritten by the authors. The author(s) reviewed and edited the output as needed and take full responsibility for the content of the published article.

## Data Availability Statement

All experimental data are available upon request

## Acknowledgments

This work was funded by the author’s employers, Repligen, etherna, and Recipharm, and Refeyn.

## Author Contributions

NEC, HM, CJN, MRS, RAW, JF, BK, MJR, LD, and SD performed experiments and analyzed data. NEC prepared figures and drafted the manuscript. All authors designed experiments and edited the manuscript.

## Declaration of interest statement

All authors are current or former employees of the companies stated in the affiliation section. NEC and TCS are co-inventors on a patent related to dsRNA affinity purification resins.

---

**Supplementary Figure 2.**
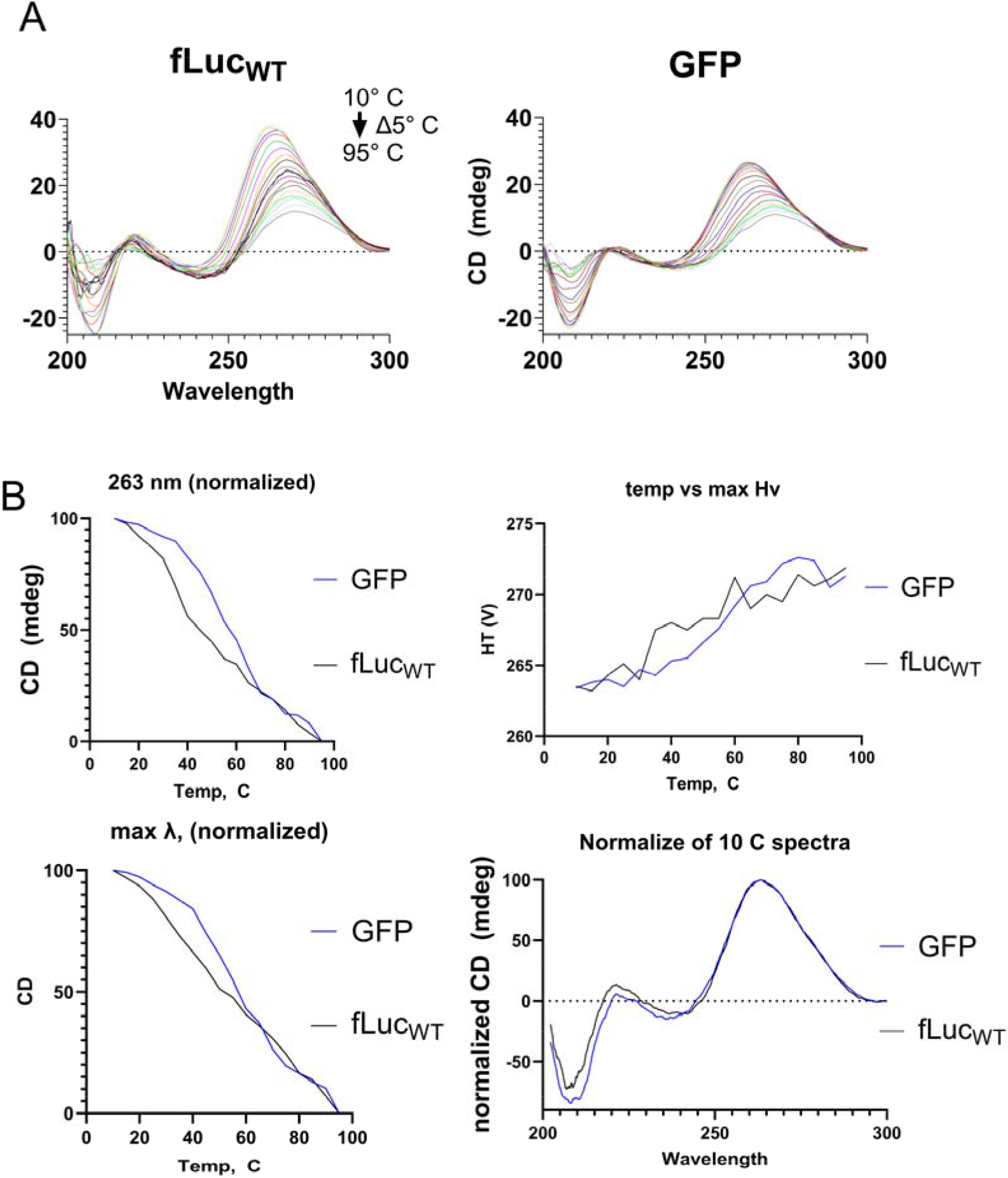
Circular dichroism of 4 mRNA samples. (A) Spectra were collected from 200 to 300 nm at temperatures from 10° C to 95° C, in 5° C increments (B) Analysis of the normalized spectra Absent large differences in secondary structure, we used analytical size-exclusion chromatography to measure the differences in higher-order tertiary or quaternary structures. RNAs were incubated either in water or 0.63 M NaCl prior to injection on a 1,000 Å pore size exclusion column. For GFP in water, a large monomer peak and a minor dimer peak (Figure 3A, blue traces) were observed. For fLuc_WT_ in water, a broad distribution dominated by dimer and higher-order species was observed. After incubation in 0.63 M NaCl buffer, fLuc_WT_ shifted further toward higher-order structures (Figure 3A, red traces). In contrast, GFP was unaffected by the incubation in the high-salt buffer.

## Notes

### Summary of Updates

THe labels were cut-off on FIgure 5

