## Supplemental Sequences for "RNA self-association limits the removal of double-stranded RNA by affinity chromatography"

**Sequences, lengths, and calculated molecular weights dasher GFP, Fluc_WT_**

1) wild-type firefly luciferase (Fluc_WT_). 1,764 nucleotides, calculated 567.94 kDa

>FlucWT

gucuagaaauaauuuuguuuaacuuuaagaaggagauauaaccaugaaaaucgaagaagguaaaggucaccaucaccaucaccacggauccauggaagacgccaaaaacauaaagaaaggcccggcgccauucuauccucuagaggauggaaccgcuggagagcaacugcauaaggcuaugaagagauacgcccugguuccuggaacaauugcuuuuacagaugcacauaucgaggugaacaucacguacgcggaauacuucgaaauguccguucgguuggcagaagcuaugaaacgauaugggcugaauacaaaucacagaaucgucguaugcagugaaaacucucuucaauucuuuaugccgguguugggcgcguuauuuaucggaguugcaguugcgcccgcgaacgacauuuauaaugaacgugaauugcucaacaguaugaacauuucgcagccuaccguaguguuuguuuccaaaaagggguugcaaaaaauuuugaacgugcaaaaaaaauuaccaauaauccagaaaauuauuaucauggauucuaaaacggauuaccagggauuucagucgauguacacguucgucacaucucaucuaccucccgguuuuaaugaauacgauuuuguaccagaguccuuugaucgugacaaaacaauugcacugauaaugaauuccucuggaucuacuggguuaccuaaggguguggcccuuccgcauagaacugccugcgucagauucucgcaugccagagauccuauuuuuggcaaucaaaucauuccggauacugcgauuuuaaguguuguuccauuccaucacgguuuuggaauguuuacuacacucggauauuugauauguggauuucgagucgucuuaauguauagauuugaagaagagcuguuuuuacgaucccuucaggauuacaaaauucaaagugcguugcuaguaccaacccuauuuucauucuucgccaaaagcacucugauugacaaauacgauuuaucuaauuuacacgaaauugcuucugggggcgcaccucuuucgaaagaagucggggaagcgguugcaaaacgcuuccaucuuccagggauacgacaaggauaugggcucacugagacuacaucagcuauucugauuacacccgagggggaugauaaaccgggcgcggucgguaaaguuguuccauuuuuugaagcgaagguuguggaucuggauaccgggaaaacgcugggcguuaaucagagaggcgaauuaugugucagaggaccuaugauuauguccgguuauguaaacaauccggaagcgaccaacgccuugauugacaaggauggauggcuacauucuggagacauagcuuacugggacgaagacgaacacuucuucauaguugaccgcuugaagucuuuaauuaaauacaaaggauaucagguggcccccgcugaauuggaaucgauauuguuacaacaccccaacaucuucgacgcgggcguggcaggucuucccgacgaugacgccggugaacuucccgccgccguuguuguuuuggagcacggaaagacgaugacggaaaaagagaucguggauuacgucgccagucaaguaacaaccgcgaaaaaguugcgcggaggaguuguguuuguggacgaaguaccgaaaggucuuaccggaaaacucgacgcaagaaaaaucagagagauccucauaaaggccaagaagggcggaaaguccaaacucgaguaagguuaaccugcaggagg

4) Dasher green fluorescent protein. 711 nucleotides, calculated 228.89 kDa

(Source: https://www.snapgene.com/plasmids/fluorescent_protein_genes_and_plasmids/daGFP)

>GFP(Dasher)

Augacagcuuuaacugaaggggccaagcuuuucgaaaaagagaucccauauaucaccgaacuugaaggugacguugaggguaugaaguuuaucauaaaaggagaaggcacaggugaugcgacaacggguacaaucaaggcaaaguacauuuguacaacuggggaccugccugucccaugggccacuuuggugucuacuuugucuuacggcguacaauguuuugcuaaguacccuucacacaucaaagauuucuuuaagucugcaaugccugaaggauacacacaggaacguacaauuucauuugagggcgacggugucuauaaaacaagagcuaugguuacuuaugaaagagguuccaucuacaacagagugacacuaacgggcgaaaauuucaaaaaggauggacauauuuugcguaaaaacguagcuuuccaaugcccaccaucaauacuauacauucugccagauacuguaaacaaugguauuagagucgaguuuaaucaagcuuaugauauagaaggugucacugaaaaauugguuacaaaaugcagccaaaugaauagaccauuggcaggaucugccgcugugcauaucccuagauaccaucacauuaccuaccacaccaaauuaaguaaagacagggaugaacgaagagaucauauguguuuaguugagguuguuaaggcaguugaucucgacacuuaccaauaa
